# Pathogenic mutations in *NUBPL* affect complex I activity and cold tolerance in the yeast model *Yarrowia lipolytica*

**DOI:** 10.1101/338517

**Authors:** Andrew E. Maclean, Virginia E. Kimonis, Janneke Balk

**Affiliations:** Department of Biological Chemistry, John Innes Centre, Norwich NR4 7UH, UK; School of Biological Sciences, University of East Anglia, Norwich NR4 7TJ, UK; Division of Genetics and Metabolism, Department of Pediatrics, University of California, Irvine, <u>Children’s Hospital of Orange County</u>, Orange, CA, USA

## Abstract

Complex I deficiency is a common cause of mitochondrial disease, resulting from mutations in genes encoding structural subunits, assembly factors or defects in mitochondrial gene expression. Advances in genetic diagnostics and sequencing have led to identification of several variants in NUBPL, an assembly factor of complex I, which are potentially pathogenic. To help assign pathogenicity and learn more about the function of NUBPL, amino acid substitutions were recreated in the homologous Ind1 protein of the yeast model *Yarrowia lipolytica*. L102P destabilized the Ind1 protein, leading to a null-mutant phenotype. D103Y, L191F and G285C affected complex I assembly to varying degrees, whereas the G138D variant did not impact on complex I levels or dNADH:ubiquinone activity. Blue-native PAGE and immunolabelling of the structural subunits NUBM and NUCM revealed that all Ind1 variants accumulated a Q-module intermediate of complex I. In the D103Y variant the matrix arm intermediate was virtually absent, indicating a dominant effect. Dysfunction of Ind1, but not absence of complex I, rendered *Y. lipolytica* sensitive to cold. The Ind1 G285C variant was able to support complex I assembly at 28°C, but not at 10°C. Our results indicate that Ind1 is required for progression of assembly from the Q module to the full matrix arm. Cold sensitivity could be developed as a phenotype assay to demonstrate pathogenicity of *NUBPL* mutations and other complex I defects.

## Introduction

Defects in respiratory chain function underlie a large share of mitochondrial disorders, of which a third are due to complex I deficiency (OMIM 252010) (1–3). Clinical presentation such as Leigh syndrome usually occur in infancy or early adulthood. Symptoms include skeletal muscle myopathy, cardiomyopathy, hypotonia, stroke, ataxia and lactic acidosis (3, 4). Human complex I has 44 different structural subunits, of which 37 are encoded in the nuclear genome and 7 in the mitochondrial genome (5). Disease-causing mutations have been identified in 26 out of 44 structural genes. However, this only provides a diagnosis for about 50% of cases, as complex I deficiency may also be caused by mutations in assembly factors or mitochondrial gene expression (2, 6). Assembly factors are defined as proteins which are required for the correct assembly and function of complex I but are not present in the mature structure. So far, 16 assembly proteins have been identified for complex I, and disease-causing mutations have been found in 10 of those (3, 7). Particularly, advances in genome sequencing have led to the identification of several novel genes affecting complex I function, providing a genetic diagnosis for patients with these rare inheritable conditions.

The complex I assembly factor NUBPL (NUcleotide Binding Protein-Like) was first identified in the aerobic yeast *Yarrowia lipolytica* where it was named Ind1 (Iron-sulfur protein required for NADH dehydrogenase) (8). Genomic deletion of *IND1* resulted in a major decrease in complex I, to approximately 28% of wild-type levels, but it has no effect on other respiratory complexes. Ind1 has been proposed to insert iron-sulfur (FeS) clusters in complex I based on its homology to the cytosolic FeS cluster assembly factors Nbp35 and Cfd1, corresponding to NUBP1 and NUBP2, respectively, in human (9). However, it should be noted that Ind1 is not essentially required for complex I in *Y. lipolytica* and it would therefore only have an auxiliary role in cofactor assembly.

Phylogenetic analysis showed that the *IND1* gene is present in almost all eukaryotes, and closely matches the distribution of complex I (8). A study of the human homologue showed that siRNA knockdown of *NUBPL* in HeLa cells led to a specific decrease in complex I, as well as accumulation of assembly intermediates (10). Moreover, mutations in *NUBPL* were identified in patients with complex I deficiency by exome sequencing (1, 11, 12). Recently, *NUBPL* was found among 191 genes that are essential for oxidative phosphorylation in a genome-wide screen of CRISPR mutants (13). The *IND1* homologue in the model plant *Arabidopsis thaliana* is also required for complex I assembly (14). Combined, these studies show that NUBPL has an evolutionary conserved function in complex I assembly, but its precise molecular function remains to be demonstrated.

So far, 12 different cases of mutations in *NUBPL* have been reported or newly diagnosed (Table 1). Patients display various symptoms within the broad spectrum of mitochondrial disorders, including motor problems, increased lactate levels and some degree of complex I deficiency. In addition, NUBPL patients show a very distinct magnetic resonance imaging (MRI) pattern with abnormalities of the cerebellar cortex, deep cerebral white matter and corpus callosum (11). Genetic diagnoses by exome sequencing revealed that all patients have inherited a branch-site mutation in *NUBPL*, c.815-27T>C, from either the mother or father. Except for one suspected homozygous case (Table 1, patient 3), all other cases are compound heterozygous carrying a likely deleterious mutation in the other copy of the *NUBPL* gene (Table 1). The c.815-27T>C mutation affects a splice recognition site in intron 9, leading to aberrant mRNA splicing (15). Approximately 30% of wild-type *NUBPL* transcript remains but two miss-spliced transcripts are also produced. One of these disappears by nonsense mediated decay. The other miss-spliced transcript lacks exon 10, leading to D273Q, a frameshift and then a premature stop codon. The predicted protein product, p.D273QfsX31, is unstable, as NUBPL protein is almost undetectable in patient fibroblasts (15). c.815-27T>C is often co-inherited with a missense c.G166>A mutation (p.G56R), which, on its own, is not thought to be pathogenic (15), and absent in at least two cases (Table 1, patient 2 and siblings 9 and 10).

**Table 1.** Overview of mutations in *NUBPL* (NG_028349.1) associated with complex I deficiency or mitochondrial disease (OMIM 252010).

| Patient | cDNA | Protein | Site | Paternal or Maternal | Reference |
| --- | --- | --- | --- | --- | --- |
| 1 | c.166G>A | p.G56R | Exon 2 | Paternal | (1, 15) |
|  | c.815-27T>C<sup>a</sup> | p.D273QfsX31 | Intron 9 | Paternal |  |
|  | 240-kb deletion |  | Exons 1-4 | Maternal |  |
|  | 137-kb duplication |  | Exon 7 | Maternal |  |
| 2 | c.815-27T>C | p.D273QfsX31 | Intron 9 | Paternal | (12) |
|  | c.205_206delGT | p.V69Yfs*80 |  | Maternal |  |
| 3<sup>b</sup> | c.166G>A | p.G56R | Exon 2 | N/A | (11) |
|  | c.815-27T>C | p.D273QfsX31 | Intron 9 | N/A |  |
| 4 | c.166G>A | p.G56R | Exon 2 | Paternal | (11) |
|  | c.815-27T>C | p.D273QfsX31 | Intron 9 | Paternal |  |
|  | c.667_668insCCT TGTGCTG | p.E223Afs*4 | Exon 8 | Maternal |  |
| 5,6 | c.166G>A | p.G56R | Exon 2 | Paternal | (11) |
|  | c.815-27T>C | p.D273QfsX31 | Intron 9 | Paternal |  |
|  | c.313G>T | p.D105Y | Exon 4 | Maternal |  |
| 7 | c.166G>A | p.G56R | Exon 2 | Paternal | (11) |
|  | c.815-27T>C | p.D273QfsX31 | Intron 9 | Paternal |  |
|  | c.693+1G>A | p.? | Intron 8 | Maternal |  |
| 8 | c.579A>C | p.L193F | Exon 7 | Paternal | (11) |
|  | c.166G>A | p.G56R | Exon 2 | Maternal |  |
|  | c.815-27T>C | p.D273QfsX31 | Intron 9 | Maternal |  |
| 9, 10 | c.815-27T>C | p.D273QfsX31 | Intron 9 | Paternal | Virginia Kimonis, *unpublished* |
|  | c.311T>C | p.L104P | Exon 4 | Maternal |  |
| 11 | c.166G>A | p.G56R | Exon 2 | N/A | Holger Prokisch, Helmholtz Zentrum München (*pers. communication*) |
|  | c.815-27T>C | p.D273QfsX31 | Intron 9 | N/A |  |
|  | c.859G>T | p.G287C | Exon 10 | N/A |  |
| 12 | c.815-27T>C | p.D273QfsX31 | Intron 9 | Maternal | Virginia Kimonis, *unpublished* |
|  | c.166G>A | p.G56R | Exon 2 | Maternal |  |
|  | c.311T>C | p.L104P | Exon 4 | Paternal |  |

<sup>a</sup> The paternal c.166G>A mutation and the maternal exon deletions/duplication were identified by exome sequencing (1), but further analysis identified the c.815-27T>C splice site mutation in the paternal allele (15).
<sup>b</sup> Kevelam et al (Ref. (11)) could not confirm whether the mutations were homozygous or hemizygous because no parental DNA or fibroblasts were available for study. However, given the other cases, it is possible that another mutation in *NUBPL* is present to give a compound heterozygous situation.

The haploid yeast *Y. lipolytica* has been used extensively in biotechnology but it has also been developed as a model organism to study complex I biology (16–20). This has been necessary because the usual yeast model Baker’s yeast, *Saccharomyces cerevisae*, does not have complex I. The D273QfsX31 variation of NUBPL was recreated in the Ind1 protein of *Y. lipolytica*, which confirmed that the altered protein was unable to support the assembly of complex I (21). Here, we compare six mutations in the *NUBPL* gene, 3 of which are newly identified, in an effort to assign pathogenicity and to enhance our understanding of the molecular function of NUBPL/Ind1. The resulting protein variants were tested in *Y. lipolytica* for protein stability and complex I levels, oxidoreductase activity and assembly intermediates. Together, the results indicate pathogenicity of four missense mutations and reveal that *ind1* mutants are sensitive to low temperature.

## Results

### Selection of *NUBPL* mutations for analysis in *Y. lipolytica*

Twelve cases of mitochondrial disorders associated with mutations in *NUBPL* have been published or were otherwise known to us (Table 1). In addition, many more polymorphisms in *NUBPL* have been uncovered by large-scale exome sequencing (exac.broadinstitute.org), of which V182A and G138D occur at a high allele frequency (Table 2). Mutations that result in early frameshifts, insertions or deletions (patients 1, 2, 4 and 7) are almost certainly deleterious but are unlikely to shed light on the molecular function of NUBPL. Mutations resulting in a single amino acid change at a conserved position, such as L104P (patients 9,10 and 12), D105Y (patients 5,6), G138D (ExAC database), L193F (patient 8) and G287C (patient 11) were chosen for functional analysis in *Y. lipolytica*. V182 is only semi-conserved in *Y. lipolytica* Ind1, corresponding to M180, and was therefore not included in this study.

**Table 2.** Allele frequencies of NUBPL variants.

| Variant | Allele Count | Total Alleles | Allele Frequency | Conserved | PolyPhen-2 |
| --- | --- | --- | --- | --- | --- |
| N198T | 529 | 119894 | $4.41 \times 10^{-3}$ | No | 0.888 |
| c.815-27T>C | 436 | 117040 | $3.73 \times 10^{-3}$ | N/A | N/A |
| V182A | 348 | 120522 | $2.89 \times 10^{-3}$ | No | 0.923 |
| G26V | 124 | 82280 | $1.51 \times 10^{-3}$ | No | 0.041 |
| H229Y | 137 | 120644 | $1.14 \times 10^{-3}$ | No | 0.754 |
| G138D | 88 | 117184 | $7.51 \times 10^{-4}$ | Yes | 0.999 |
| S128N | 25 | 116542 | $2.13 \times 10^{-4}$ | No | 0.000 |
| L104P | 25 | 120170 | $2.08 \times 10^{-4}$ | Yes | 1.000 |
| D105Y | 1 | 120200 | $8.319 \times 10^{-6}$ | Yes | 1.000 |
| L193F | N/A | N/A | N/A | Yes | 1.000 |
| G287C | N/A | N/A | N/A | Yes | 1.000 |

Allele frequency data from ExAC (http://exac.broadinstitute.org/). The top 9 variants in conserved amino acids are listed, ranked by allele frequency, as well as clinically relevant variants (c.815-27T>C, D105Y, L193F, G287C). A PolyPhen-2 score close to 1 indicates that the amino acid change is “probably damaging”.

L104P and D105Y affect residues in the highly conserved Switch I motif (Figure 1A) involved in ATP hydrolysis; L193F affects a residue just outside the Mrp family signature; and G287C introduces an additional cysteine to the C-terminus. The corresponding amino acids in the Ind1 protein of *Y. lipolytica* were identified by alignment using Clustal Omega (Figure 1A, B) and this amino acid numbering is used throughout the study. A homology model of Ind1 was made and the positions of the substituted amino acids are indicated in Figure 1C.

**Figure 1.**
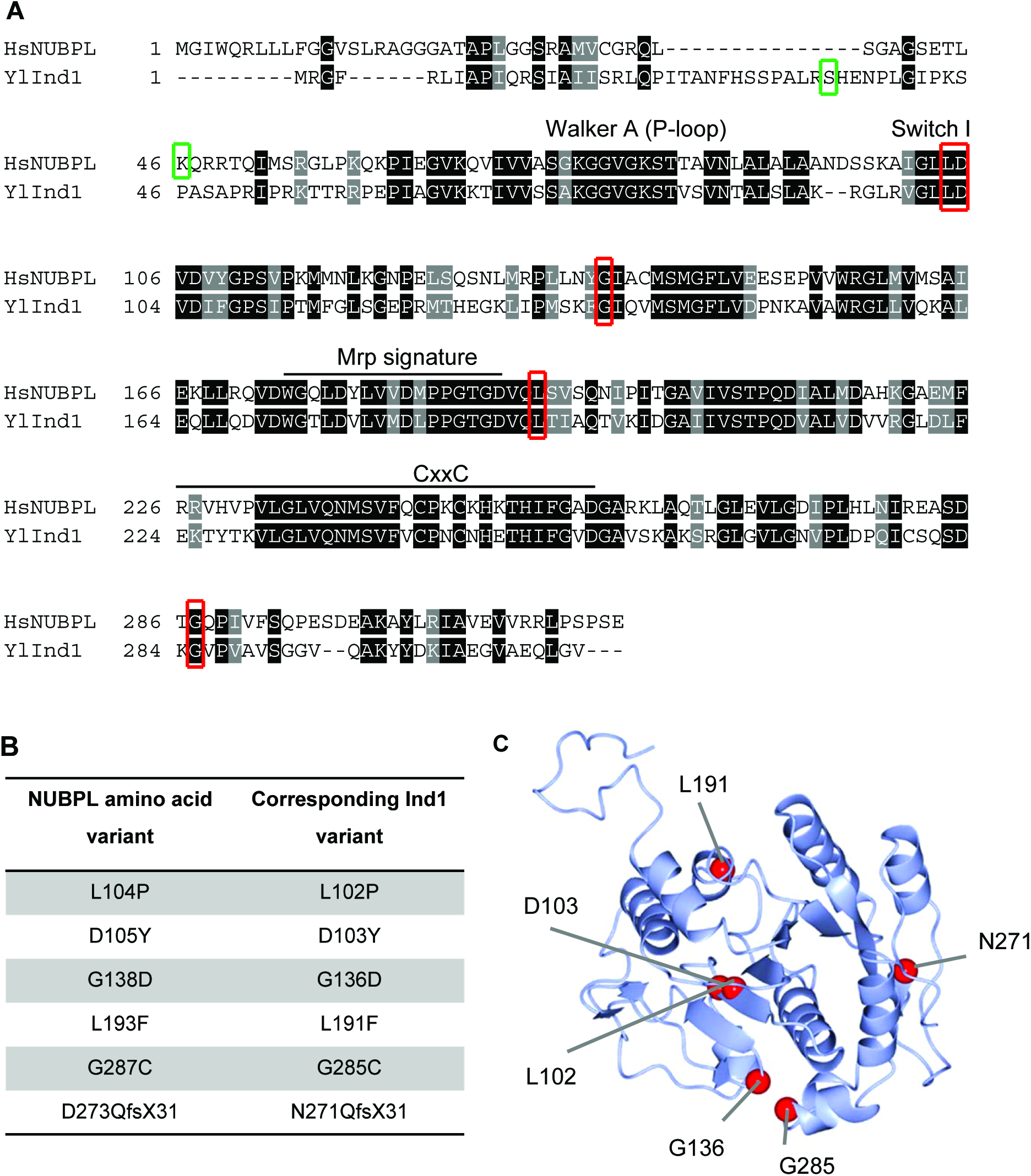
Amino acid substitutions in NUBPL and their position in the homologous Ind1 protein in *Y. lipolytica*. (**A**) Alignment of amino acid sequences of human (Hs) NUBPL (NP_079428.2) and *Y. lipolytica* (Yl) Ind1(XP_501064.1) using Clustal Omega and Boxshade for colouring. Amino acid changes that are investigated in this study are indicated with red boxes. The start of the mature N-terminus is indicated with green boxes. The Walker A motif (P-loop), Switch I, Mrp family signature (ProSite entry PS01215) and CxxC motif are indicated. (**B**) Protein variants in human NUBPL and the corresponding amino acid changes in *Y. lipolytica* Ind1 based on alignment. (**C**) Homology model of Ind1 using IntFOLD server using template 3vx3A.pdb (HypB from *Thermococcus kodakarensis* KOD1). Amino acid residues that were substituted in this study are indicated with red spheres.

### Protein stability in Ind1 variants

In order to study the biochemical and physiological effects of the amino acid changes, site-directed mutations were introduced into a plasmid carrying the *IND1* gene under the control of its native promoter and transformed into *Y. lipolytica ind1Δ* cells, a GB10 strain with a genomic deletion of the *IND1* gene (8). The plasmid contains a chromosomal ARS/CEN region which is maintained at ~1 copy per cell. The GB10 strain was engineered to contain the *NDH2i* gene, encoding an alternative NADH dehydrogenase targeted to the matrix side of the inner mitochondrial membrane, which bypasses the essential requirement of respiratory complex I in *Y. lipolytica* (22). The previously reported variant Ind1 protein N271QfsX31, recapitulating the effect of the c.815-27T>C branch-site mutation in *NUBPL*, was included for comparison (21).

Mitochondrial membranes of each strain were isolated and subjected to Western blot analysis to visualize Ind1 protein levels. The D103Y, G136D, L191F and G285C variants displayed levels of Ind1 protein similar to *ind1*Δ + *IND1* (henceforth referred to as complemented wild type, cWT) (Figure 2A). The L102P substitution resulted in lower levels of Ind1 protein, to approximately 45% of cWT levels. The decrease in L102P protein was similar to that seen in N271QfsX31. Antibodies against aconitase (Aco1) and subunit 2 of complex II (Sdh2) were used as a loading control and to confirm that other FeS-cluster binding proteins are not affected (Figure 2A).

**Figure 2.**
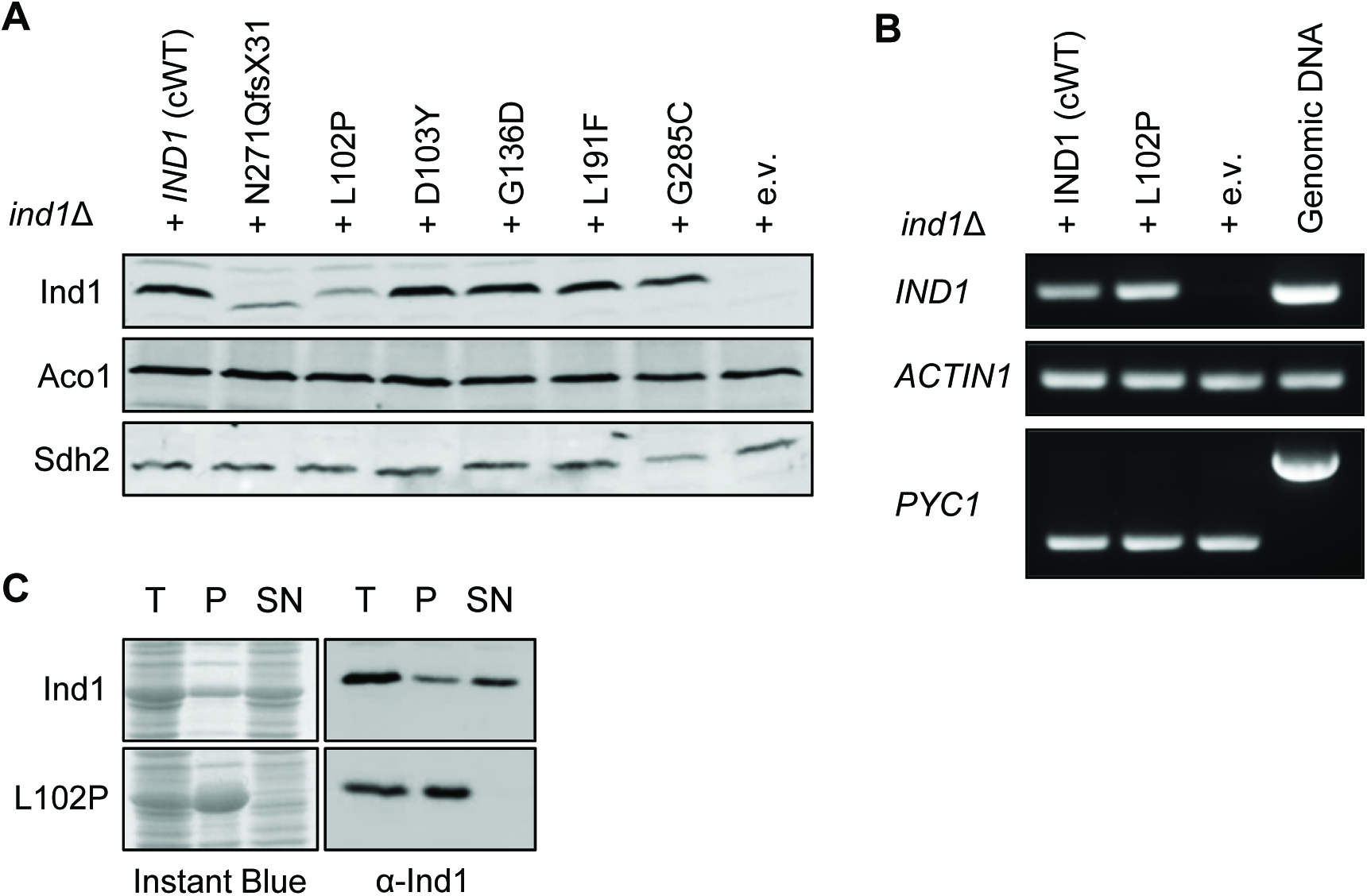
Expression of variant Ind1 proteins in *Y. lipolytica*. (**A**) Protein blot analysis of Ind1 in mitochondrial membranes from strains expressing wild-type *IND1* (cWT) and the indicated protein variants, in the *ind1*Δ background. The N271QfsX31 protein is of lower molecular weight due to truncation of the C-terminus by 11 amino acids. Antibodies against the mitochondrial protein aconitase (Aco1) and subunit 2 of succinate dehydrogenase (Sdh2) were used to confirm protein equal loading and stability of these FeS cluster proteins in *ind1* mutants. (**B**) Transcript levels of IND1^L102P^ compared to wild-type *IND1* by semi-quantitative RT-PCR. A PCR reaction for *PYC1 (YALI0C24101g*), containing an intron, showed that the cDNA samples were free from genomic DNA and equal in cDNA content. *ACTIN1* was used as an additional control for the amount of cDNA template. (**C**) Solubility of wild-type Ind1 and Ind1^L102P^ protein expressed in *E. coli*. Total protein extract (T) was sonicated and centrifuged to give insoluble pellet (P) and soluble supernatant (SN). Gels were stained with Instant Blue (left panels) or immunolabelled for Ind1 (right panels).

To rule out that lower amounts of the Ind1^L102P^ protein were due to decreased transcript levels, expression of *IND1* was assessed by RT-PCR (Figure 2B). Normal levels of *IND1* transcript in the L102P variant indicated that the protein undergoes post-translational degradation. In order to further investigate the effect of L102P on protein stability, Ind1 and Ind1^L102P^ were expressed in *E. coli*. Separation of soluble and insoluble fractions showed that all Ind1^L102P^ protein is found in the insoluble pellet fraction, whilst most of the normal Ind1 protein can be found in the soluble fraction. This suggests that the L102P substitution causes protein misfolding, which would result in its degradation by proteases. Overall these data show that, with the exception of L102P, the selected amino acid substitutions in Ind1 do not affect protein stability.

### Ind1 variants L102P, D103Y and L191F have strongly decreased levels of complex I

It has previously been shown that cells lacking *IND1* have approximately 28% of fully active complex I compared to wild type (8). To investigate the effect of the amino acid variants in Ind1 on complex I, mitochondrial membranes were separated by blue-native PAGE (BN-PAGE) to resolve the respiratory complexes. Incubation of the gel with NADH and nitro-blue tetrazolium (NBT) reveals NADH dehydrogenase activities, which is used as a proxy to estimate complex I levels (23). In cells expressing the Ind1 variants L102P and D103Y, complex I was strongly decreased, similar to the levels in the *ind1*Δ mutant (empty vector control, e.v.) and N271QfsX31 (Figure 3A). The L191F substitution resulted in a minor decrease in complex I, whereas G138D and G285C were indistinguishable from cWT. For comparison, a knockout strain of the NUBM subunit of complex I (*nubm*Δ) displayed no complex I activity, in agreement with the FMN cofactor of NUBM being the site of NADH oxidation. Another complex I mutant, where a key catalytic residue of the NUCM subunit has been substituted (Y144F) (24), displayed complex I levels similar to cWT. This is consistent with a mutation that affects catalysis but not assembly of complex I. Complex V levels were unaffected by mutations in *IND1* (Figure 3A, lower Coomassie-stained band).

**Figure 3.**
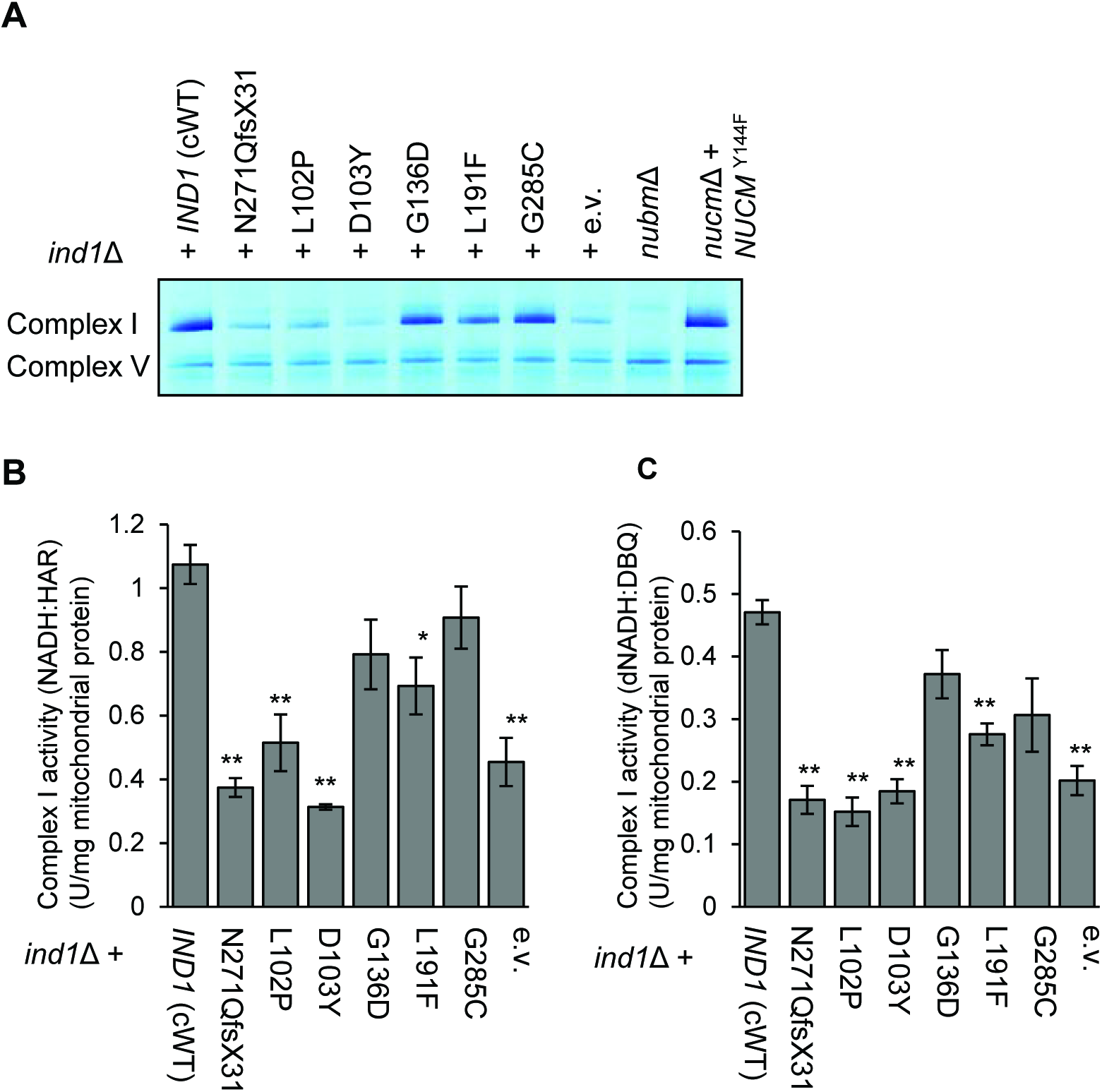
Complex I levels and oxidoreductase activities in Ind1 variants. (**A**) Complex I levels shown by in-gel NADH:NBT staining of mitochondrial membranes from *ind1*Δ producing wild-type *IND1* (cWT) and the indicated protein variants, compared to two characterized complex I mutants (*nubm*Δ and *nucm*Δ + *NUCM*^Y144F^). (**B**) NADH:HAR and (**C**) dNADH:DBQ oxidoreductase activity in mitochondrial membranes from the indicated strains. Bars represent the mean ± SE (n = 3 biological replicates). * p<0.05, ** p<0.01 (two-sample *t*-test). Numerical values are given in Table S1.

To measure oxidoreductase activity of the matrix arm of complex I, two different spectrophotometric assays were used. Electron transfer from NADH to the artificial electron acceptor hexaammineruthenium(III) chloride (HAR) involves only the primary electron acceptor FMN bound to the NUBM subunit of complex I and, like NADH:NBT staining, serves as a proxy for complex I content. The L102P, D103Y, L191F, and N271QfsX31 variants showed a significant decrease in NADH:HAR activity, whereas G136D and G285C were similar to cWT (Figure 3B, Table S1).

To assess NADH:ubiquinone oxidoreductase activity, electron transfer from deamino-NADH (dNADH) to n-decylubiquinone (DBQ) was measured, encompassing all cofactors in the matrix arm of complex I. A significant decrease in electron transfer was seen with the L102P, D103Y, L191F, and N271QfsX31 variants, but not in G136D and G285C (Figure 3C, Table S1). The *nucmΔ* + *NUCM*^Y144F^ mutant had low dNADH:DBQ activity (Table S1), despite wildtype levels of fully assembled complex I (Figure 3A). Comparison of dNADH: DBQ activity between the Ind1 variants and *nucm*Δ + *NUCM*^Y144F^ shows that in the Ind1 variants there is still substantial activity remaining. For the Ind1 variants, the NADH:HAR activities correspond with those of dNADH:DBQ, suggesting that all the complex I present is enzymatically active. Taken together, the data from three different methods are in agreement and indicate that the L102P and D103Y substitutions in Ind1 cause a defect in complex I assembly of similar severity as *IND1* deletion. The L191F substitution caused a mild decrease, whereas G138D and G285C appeared to have no significant effect on complex I levels under standard growth conditions.

### All *ind1* mutants accumulate a Q module assembly intermediate

In human cell lines depleted of *NUBPL*, a subcomplex representing part of the membrane arm was observed (10). Similarly, the Arabidopsis *indh* mutant had a membrane arm assembly intermediate, but no full-size complex I (14). To investigate complex I assembly in *Y. lipolytica* cells lacking Ind1, the *ind1*Δ strain was compared to mutant strains of different subunits of the matrix arm of complex I, *nubm*Δ, *nucm*Δ and *nukm*Δ. NUBM is the homologue of the human NDUFV1 protein (bovine 51-kDa subunit) in the N-module. NUCM (human NDUFS2, bovine 49 kDa) and NUKM (human NDUFS7, bovine PSST) are in the Q-module (Figure 4A). In the *ind1*Δ mutant, the protein levels of NUBM are slightly decreased but the levels of NUCM are not affected (Figure 4B). Thus, compared to the decreased levels of complex I in *ind1*Δ, NUBM and NUCM are imported into the mitochondrial matrix but the majority is not assembled. Decreased levels of NUBM are also seen in the *nucm*Δ and *nukm*Δ mutants, indicating that if the Q-module is not assembled, NUBM is degraded.

**Figure 4.**
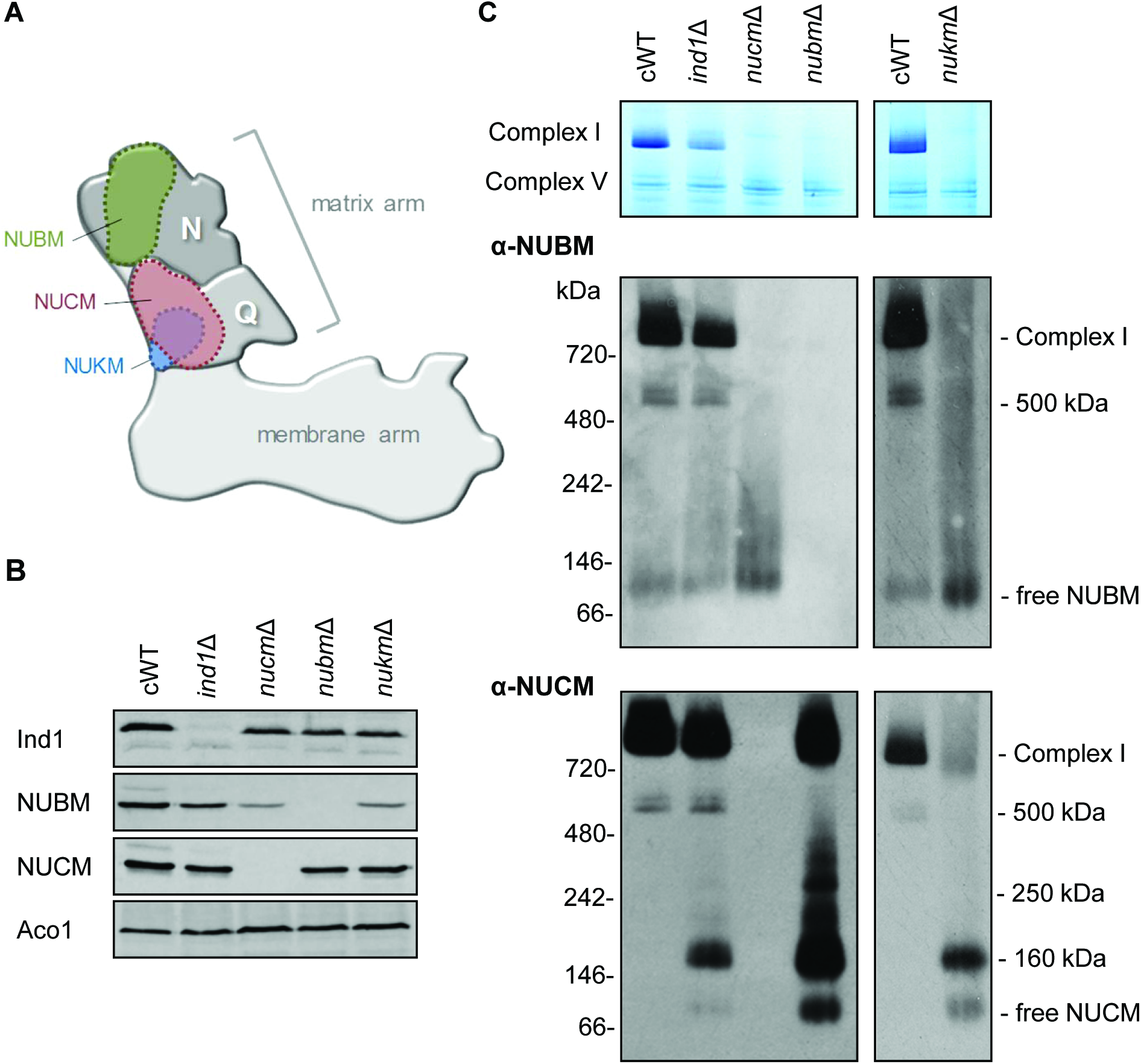
Complex I assembly in *ind1*Δ compared to deletion mutants of other subunits. (**A**) Diagram of complex I showing the position of NUBM in the N-module and NUCM, NUKM in the Q-module of the matrix arm. (**B, C**) Mitochondrial membranes from the indicated complex I strains (complemented wild type cWT, *ind1*Δ, *nucm*Δ, *nubm*Δ and *nukm*Δ) were separated by SDS-PAGE (B) or by BN-PAGE (C). Western blots were labelled with antibodies against Ind1 and the complex I subunits NUBM and NUCM, as indicated. Antibodies against mitochondrial aconitase, Aco1, were used to confirm equal loading and protein transfer. The immunoblots were exposed for a relatively long time to visualize all assembly intermediates, which also revealed residual complex I lacking NUBM in the *nubm*Δ mutant.

Mitochondrial membranes were isolated from cWT, *ind1*Δ, *nubm*Δ, *nucm*Δ and *nukm*Δ and the respiratory complexes were separated by BN-PAGE, followed by Western blotting with antibodies against NUBM and NUCM. This confirmed the lack of complex I in the *nucm*Δ and *nukm*Δ mutants (Figure 4C). In the *nubm*Δ mutant a nearly full-size complex I was detected by immunolabelling which has no NADH:NBT activity, consistent with NUBM being the site of NADH oxidation. In the *ind1*Δ mutant, an assembly intermediate containing both NUBM and NUCM was seen at approximately 500 kDa which was also found in cWT (Figure 4C) and is likely to represent the full matrix arm of complex I (25). An additional assembly intermediate containing NUCM, but not NUBM, was found at approximately 160 kDa. This intermediate can also be seen in *nubm*Δ and *nukm*Δ, but not in cWT and likely corresponds to the Q module. Kmita et al., (Ref 24) have previously reported a Q module assembly intermediate in *Y. lipolytica* containing the NUCM, NUGM, NUKM and NUFM subunits. Combined these subunits would have a molecular weight of 116.42 kDa (calculated based on molecular weights in Ref. (27)). Assembly factors would likely also be bound to this structure, resulting in the observed molecular weight of ~160 kDa. These data show that while mitochondrial membranes of *ind1*Δ cells have some fully assembled complex I and a 500 kDa intermediate, there is accumulation of a Q-module intermediate.

Next, we investigated the assembly of complex I in the Ind1 protein variants. Interestingly, the G136D and G285C variants that have normal complex I levels (Figure 3A), visibly accumulated the Q-module intermediate containing NUCM (Figure 5B, lower panel). Generally, the pattern of assembly intermediates containing NUCM is similar in all mutants, except Ind1^D103Y^ accumulated relatively more Q-module intermediate and very little of the 500-kDa intermediate, similar to *nubm*Δ. For intermediates containing the NUBM subunit, variants with low levels of complex I (N271QfsX31, L102P and D103Y) closely resembled the empty vector control, whereas those with near normal complex I levels (G136D, L191F, G285C) more closely resemble cWT (Figure 5B, upper panel).

**Figure 5.**
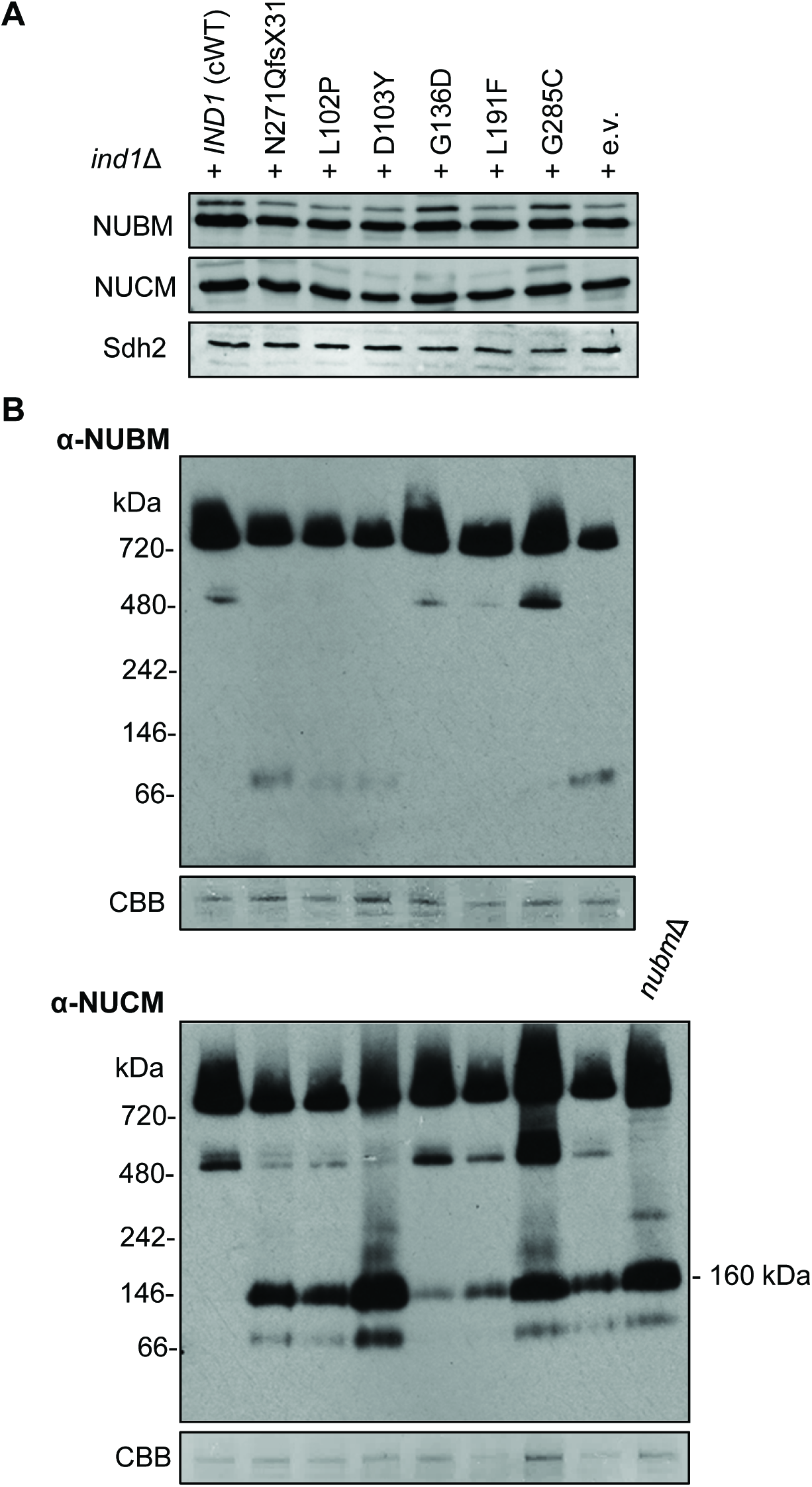
Complex I assembly intermediates in Ind1 variants. Mitochondrial membranes from *ind1*Δ cells expressing wild-type *IND1* (cWT) and the indicated protein variants were separated by SDS-PAGE (**A**) or by BN-PAGE (**B**). The gels were blotted and labelled with antibodies against NUBM, NUCM or Sdh2. CBB, Coomassie staining of complex V was used as a loading control.

### *ind1* mutants are sensitive to cold

During storage of *Y. lipolytica* strains, it was noticed that the *ind1*Δ mutant was sensitive to cold, exacerbating the mild growth defect. To investigate this further, the growth of *ind1* mutants, as well as *nucm*Δ + *NUCM*^Y144F^, *nubm*Δ and *nukm*Δ, was compared at 28°C and 10°C. Cells were spotted onto agar plates in a dilution series and, after incubation of the plates overnight at 28°C plates to initiate growth, were placed either at 28°C for a further two days or moved to 10°C for 10-14 days. At 28°C all strains grew similarly, despite slight variations in colony size (Figure 6, left panels). However, when grown at 10°C some strains displayed a dramatic growth retardation (Figure 6, right panels). The *ind1*Δ strain expressing wild-type *IND1*, Ind1 variant G136D, *nubm*Δ and *nukm*Δ all displayed significant growth at 10°C. However, all other strains at 10°C displayed very slow, or no further growth. Interestingly, the *nubm*Δ and *nukm*Δ mutants were able to grow in cold conditions but *nucm*Δ + *NUCM*^Y144F^ could not grow, similar to the *ind1*Δ mutant.

**Figure 6.**
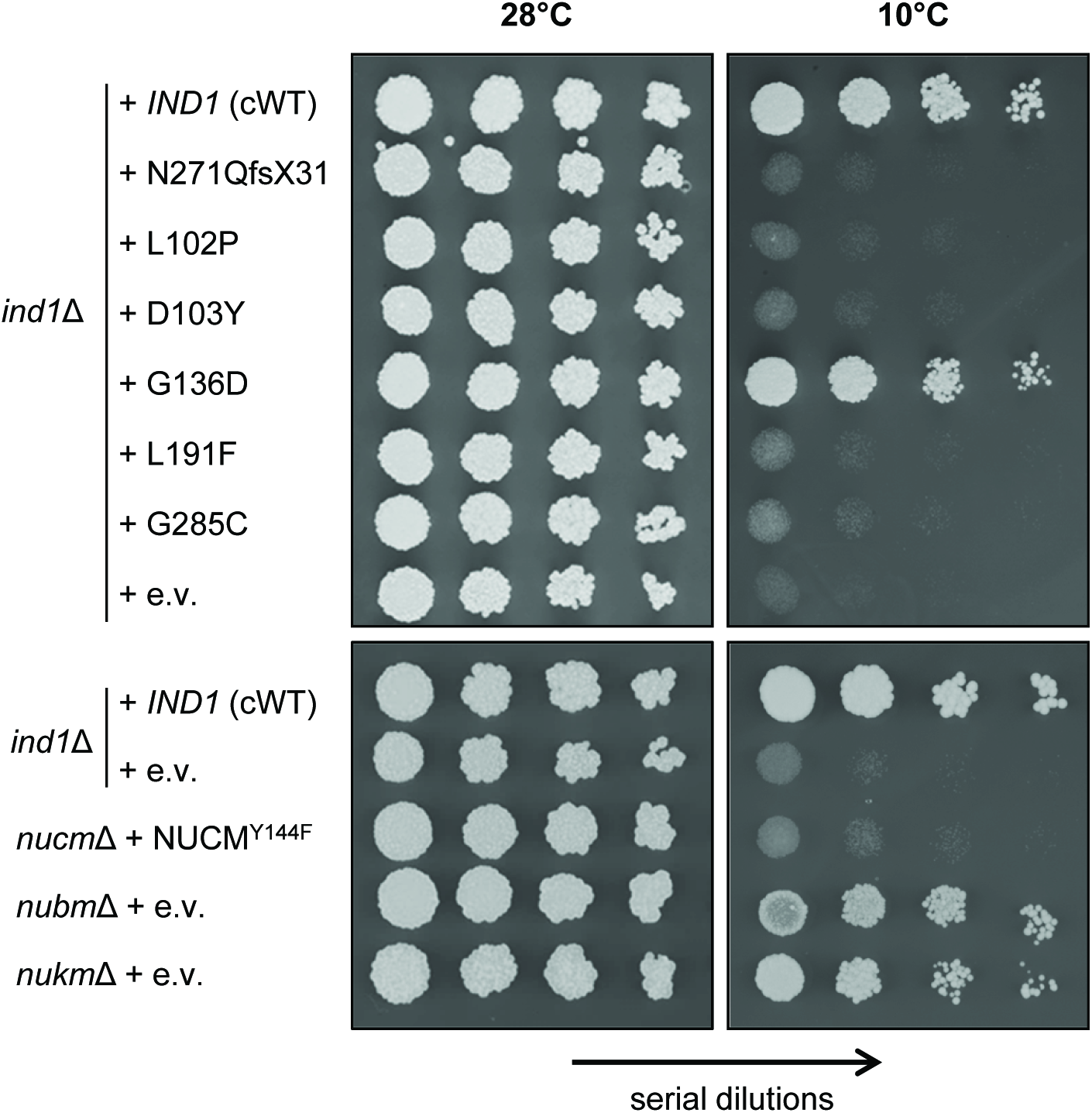
*ind1* mutants are sensitive to low temperature. Growth of *ind1*Δ cells expressing wild-type *IND1* (cWT) and the indicated protein variants, and complex I mutants *nucm*Δ + *NUCM*^Y144F^, *nubm*Δ and *nukm*Δ grown at normal (28°C) and cold (10°C) temperatures. e.v., empty vector. Images show serial dilutions of cultures spotted onto agar plates.

These data reveal a novel growth difference of certain complex I mutants grown at normal (28°C) or cold (10°C) conditions. Ind1 variants, except for G136D, also display this temperature-dependent growth defect.

### Complex I assembly in the Ind1^G285C^ variant is temperature-sensitive

The Ind1 variant G285C has normal levels and activity of complex I (Figure 3A). However, it accumulated the Q-module intermediate (Figure 5B) and growth at 10°C was impaired (Figure 6). To investigate if assembly of complex I in the G285C variant is conditional on temperature, mitochondrial membranes were isolated cells producing wild type Ind1 (cWT) and the Ind1 G285C variant after overnight growth at 10°C and compared to samples grown at 28°C. Mitochondrial membranes were separated by BN-PAGE and stained with NADH:NBT to assess formation of complex I. cWT displayed full complex I formation in cultures grown at both 28°C and 10°C (Figure 7A). By contrast, the G285C Ind1 variant could not support complex I assembly when grown at 10°C, with complex I levels decreased to those in the *ind1*Δ mutant (Figure 4C). BN-PAGE followed by Western blotting and labelling with antibodies against NUBM confirmed the significant decrease in complex I (Figure 7A). Immunodetection of Ind1 showed that the G285C protein variant is stable at 10°C, meaning the decrease in complex I levels is not due to a cold-dependent degradation of the Ind1^G285C^ protein.

**Figure 7.**
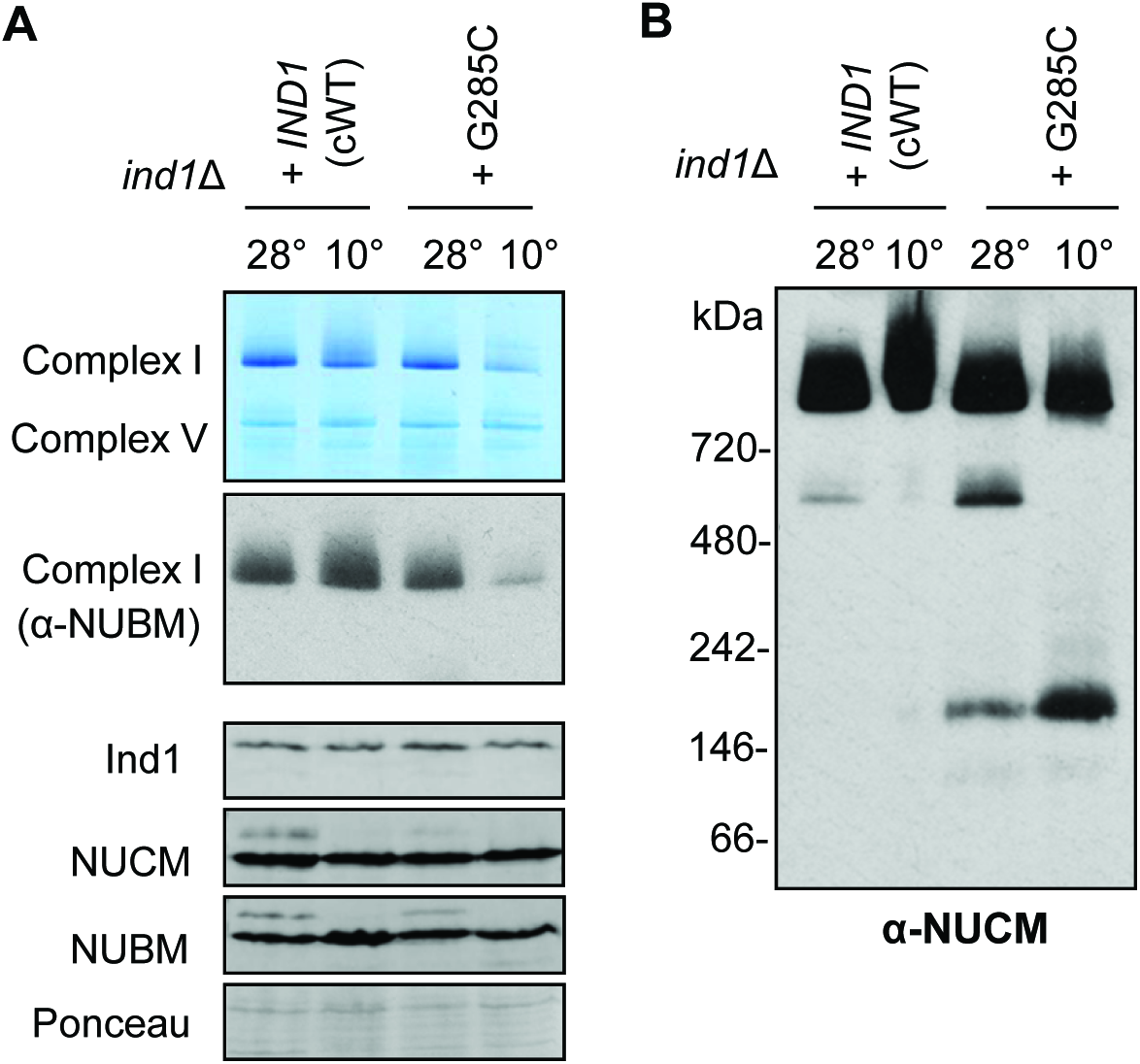
The Ind1 G285C variant cannot assemble complex I at low temperature. (**A**) Complex I levels in mitochondrial membranes from *ind1*Δ expressing wild-type *IND1* (cWT) or Ind1 variant G285C following growth at 28°C or 10°C. Complex I was visualised by BN-PAGE and NADH:NBT staining and by Western blotting with antibodies against NUBM. The levels of Ind1, NUCM and NUBM were assessed by standard SDS-PAGE and Western blot analysis. Ponceau S staining was used to confirm equal loading and transfer. (**B**) Complex I assembly intermediates containing the NUCM subunit from samples as in (A) were detected by Western blot analysis of BN-PAGE gels.

Next, the effect of cold conditions on assembly intermediates of complex I was assessed. BN-PAGE followed by Western blotting with antibodies against NUCM displayed different patterns between mitochondria isolated at normal and cold temperatures (Figure 7B). Ind1^G285C^ samples from cold conditions lacked a band corresponding to the assembly intermediate at 500 kDa, made up of the N and Q modules, and an increase in the Q module intermediate at 160 kDa. This suggests that the Ind1^G285C^ variant accumulates greater levels of the Q module at cold temperatures as it is unable to assemble it into the 500 kDa intermediate. There is still a strong signal at the size corresponding to complex I, however this is due to a long exposure time needed to see assembly intermediates. Taken together these data show that the Ind1 G285C variant is more severely impaired in complex I assembly when grown in cold conditions.

## Discussion

Since *NUBPL* was associated with mitochondrial disease resulting from complex I deficiency (1, 11), the gene is routinely included in the list of candidate genes when making a genetic diagnosis. In children’s hospitals around the world, additional mutations have been found in *NUBPL* that are likely to be pathogenic. In this study 5 protein variants of human NUBPL were recreated in the homologous Ind1 protein in the yeast *Y. lipolytica* to study the effect of the amino acid changes on complex I assembly and to confirm pathogenicity.

Ind1 variants L102P and D103Y showed the strongest decrease in complex I, similar to the effect of N271QfsX31 and deletion of the *IND1* gene (Figure 3). The L102P substitution destabilized the Ind1 protein (Figure 2), which most likely explains the complete loss of functionality. The D103Y substitution does not affect protein stability, suggesting that the amino acid change renders Ind1 inactive. Asp103 is situated at the N-terminal end of the Switch I motif (Figure 1A). In molecular motors driven by ATP (or GTP) hydrolysis, the Switch I and II sequences form the nucleotide binding pocket, while a conserved lysine residue of the Walker A motif interacts with ATP (28). Upon ATP hydrolysis, Switch I and II undergo large conformational changes, facilitating monomer-dimer transitions (e.g. in ParA and MinD in bacterial cell division) or interactions with other proteins (e.g. NifH in nitrogenase). D103 aligns with D38 in MinD of *Escherichia coli*, which residue is required for dimerization of MinD as well as interaction between MinD and MinC (29). NUBPL and other members of the NBP35/Mrp subfamily of P-loop NTPases also form dimers (9). Lack of dimer formation or lack of interaction with another protein, for example a subunit of complex I, is likely to fully abolish its function. Interestingly, the Ind1 D103Y variant appears to have a complete block in assembling the N-module onto the Q-module, as no matrix arm assembly intermediate was observed and high levels of Q-module accumulated (Figure 5B). Possibly, a failure to use ATP hydrolysis for conformational changes of Ind1 may have a dominant effect and trap a specific assembly intermediate.

The Ind1 L191F variant caused a moderate decrease in complex I levels and activity. This residue is close to the Mrp family signature (Figure 1A) found only in the subfamily of NBP35/Mrp P-loop NTPases. Members of this subfamily are found in archaea, bacteria and eukaryotes. The precise molecular function of this amino acid motif is not known.

The G136D substitution in Ind1 had no significant effect on complex I levels and redox activities (Figure 3), however the variant protein caused accumulation of the Q-module assembly intermediate, albeit at low levels (Figure 5). The equivalent G138D variant in NUBPL has so far not been associated with a clinical case. It has a relatively high allele frequency in Europeans, 1.15 × 10^-3^ versus 7.51 × 10^-4^ globally (Table 2). The allele frequency of c.815-27T>C is 4.95 × 10^-3^ in European populations (non-Finnish) and 1.26 × 10^-2^ in Finnish people. Thus, individuals homozygous for G138D or compound heterozygous for the splice-site mutation and G138D are likely to exist, and may have late onset symptoms linked to mild complex I deficiency, for example Parkinson Disease (30).

It is possible that in the case of ‘mild’ *IND1* mutations, effects on complex I assembly only manifest themselves under certain conditions, as for the Ind1 G285C variant. At the normal growth temperature of *Y. lipolytica* (28°C), there was no significant effect on complex I levels and activities, although accumulation of the Q-module assembly intermediate was observed (Figure 5). At low temperature (10°C), the Ind1 G285C variant showed a dramatic decrease in complex I and increased levels of the Q-module intermediate (Figure 7). Glycine 285 is located at the end of two short alpha helices (Figure 1C), and a cysteine may disrupt this structure. Alternatively, the cysteine, if exposed, may form an unwanted disulfide bridge with another cysteine, in particular the cysteines of the C××C motif, which are important for the function of NUBPL/Ind1 (8, 10). A Gly to Cys substitution in Nfu1, a mitochondrial protein involved in FeS cluster assembly that also carries a C××C motif, causes a dominant genetic effect in yeast (31) and severe mitochondrial disease in humans (32). Why the effects of G285C are exacerbated at cold temperatures remains an open question.

The cold-sensitive phenotype has not been reported in other mitochondrial mutants, to our knowledge. At the normal growth temperature of 28°C, *Y. lipolytica* containing Ind1 variants and complex I mutants grew similarly to cWT, because NADH oxidation in these strains is maintained by expression of ND2i, a single subunit NADH dehydrogenase on the matrix side. However, at 10°C, functionally compromised Ind1 variants and *nucm*Δ + *NUCM*^Y144F^ displayed almost no growth, whereas the *nubm*Δ and *nukm*Δ strains were comparable to cWT in cold tolerance (Figure 6). The reason for these different growth phenotypes remains unclear. An initial hypothesis is that strains with a non-functional complex I produce large amounts of reactive oxygen species (ROS) such as superoxide (5), which may be exacerbated in the cold. The *nukm*Δ mutant lacks complex I completely, and the *nubm*Δ lacks the FMN site where most ROS is produced. Further investigation is needed to uncover the cause of this growth defect. In summary four of the six mutations (L104P, D105Y, L193F and G285C) were classed as pathogenic and one (G138D) was classed as potentially pathogenic. Along with other studies (17, 20, 33, 34) these results reconfirm the utility of *Y. lipolytica* as a model for human mitochondrial disorders. However, care must be taken in over-interpreting the results as some effects seen in *Y. lipolytica* have not been found in humans. In human *NUBPL* RNAi HeLa lines, the levels of NDUFV1 subunit are substantially decreased (10), whereas in the *Y. lipolytica ind1*Δ mutant, the levels of the homologous NUBM protein are only slightly less than in cWT. Secondly, the Ind1 D103Y variant appears to have stable protein levels in *Y. lipolytica*, but human patients with D105Y have almost no NUBPL protein (11). Ideally all *NUBPL* mutations should also be tested in human cell lines, although *Y. lipolytica* is useful as a costeffective model to study potentially pathogenic mutations affecting complex I.

A defect in Q-module assembly in *Y. lipolytica ind1* mutants is consistent with studies of NUBPL in human cell lines (10, 11). The Q-module has 3 FeS clusters that may be specifically inserted by NUBPL. However, detailed proteomic analysis of complex I assembly intermediates did not find NUBPL associated with any subcomplexes, it only occurred as a dimer (35). Possibly, NUBPL interacts with an individual Q-module subunit or the protein interaction with the Q-module assembly intermediate is transient. The Ind1 protein variants characterized in this study, in particularly the D103Y variants, could serve as a tool to unravel exactly which step in complex I assembly is mediated by NUBPL/Ind1.

## Materials and Methods

### GenBank Accession Numbers

NUBPL, *Homo sapiens*: NG_028349.1; NP_079428.2

Ind1, *Yarrowia lipolytica:* XP_501064.1

### Yeast strains and growth

The *Y. lipolytica ind1*Δ strain in the GB10 genetic background (*ura3-302*, *ind1*::*URA3*, *leu2-270*, *lys11-23*, *NUGM-Htg2*, *NDH2i*, *MatB*) was used to analyse mutant versions of *IND1* and has been described previously (8). The *IND1* gene (*YALI0B18590g*) including its native promoter (1 kb upstream of the ATG) was reintroduced using the pUB4 plasmid (22) with a hygromycin selection marker, *HygB* from *Klebsiella pneumoniae*, to obtain the complemented wild type (cWT). The *nubm*Δ, *nucm*Δ, *nucm*Δ + *NUCM*^Y144F^ and *nukm*Δ were as previously described (17, 24, 36) but transformed with ‘empty’ pUB4 plasmid to grow the cells in the presence of hygromycin. *Y. lipolytica* cells were grown in rich medium containing 1% (w/v) yeast extract and 1% (w/v) glucose (Y½D), either in liquid culture or on solid medium with 2% (w/v) agar. Hygromycin B (75 μg ml-1) was added for selection of cells transformed with the pUB4 plasmid.

### Molecular cloning and mutagenesis

The *IND1* sequence from pUB4-*IND1-strep* (8) was cloned into the pGEM-T Easy vector (Promega) and used for mutagenesis using a Quikchange® II kit (Agilent), as per the manufacturer’s instructions. Mutagenesis primers were designed using the online QuikChange primer design tool, hosted at http://www.genomics.agilent.com/primerDesignProgram.jsp (Table S2). Mutated IND1-strep sequence was inserted into pUB4, between XbaI and NsiI restriction sites. Plasmids were confirmed by sequencing and diagnostic restriction digest.

*Y. lipolytica* cells were transformed as described (37). Briefly, cells were grown overnight in YPμD (1% (w/v) yeast extract, 2% (w/v) peptone and 1% (w/v) glucose, at 28°C, and collected by centrifugation, or alternatively harvested from a fresh YP½D plate. Cells were resuspended in 100 μl buffer (45% (w/v) polyethylene glycol 4000, 0.1 M lithium acetate pH 6.0, 0.1 M dithiothreitol, 250 μg ml^-1^ single-stranded carrier DNA) and 200-500 ng plasmid was added. The mixture was incubated at 39°C for 1 hour and then spread on a YP½D plate containing 75 μg ml-1 Hygromycin B. Transformants were visible after 2-3 days growth at 28°C.

### Recombinant protein expression

*IND1* sequences encoding the Ind1 variants L102P and D103Y, excluding the first 36 amino acids, were cloned from pUB4 into pET15b. Ind1-strep was expressed and purified as described in Bych et al., (Ref. (8)). Briefly, plasmids were transformed into *Escherichia coli* Rosetta containing a plasmid with the genes for Iron Sulfur Cluster assembly (pISC) and plasmidpLYS. Colonies were grown overnight at 37 °C in Lysogeny broth (LB) media, with required antibiotics, then diluted 50 × in 100 ml Terrific broth (47.6 g l^-1^ Teriffic Broth, 0.8 ml 50% (v/v) glycerol) and grown until OD600 = 0.6. Protein expression was induced by addition of 1 ml l^-1^ benzylalcohol, 50 μM Fe-ammonium citrate, 100 μM L-cysteine and 1.2 ml l^-1^ Isopropyl β-D-1-thiogalactopyranoside (IPTG). Cells were grown at 20 °C overnight, then harvested by centrifugation at 5000 × *g* and flash frozen in liquid nitrogen. Cells were then resuspended in buffer W (100 mM Tris-HCl, pH 8.0, 150 mM NaCl) with 0.2% (w/v) dodecyl maltoside. The cell suspension was sonicated, and separated into soluble and insoluble fractions by centrifugation at 16,100 × g.

### Blue-Native Polyacrylamide Gel Electrophoresis (BN-PAGE)

Unsealed mitochondrial membranes were isolated as published (16) with minor modifications. Briefly, *Y. lipolytica* cells from 0.5 l overnight culture were harvested by centrifugation at 3,500 × g. Pellet weights were typically between 2-8 g. Cells were washed with dH20. Per gram cells, 2g fine glass beads, 2 mM PMSF and 2 ml ice cold mito-membrane buffer (0.6 M sucrose, 20 mM MOPS-NaOH, pH7.5, 1 mM EDTA) were added. Cells were disrupted by 15 rounds of 1 min vortexing and 1 min incubation in ice. Differential centrifugation was performed at 3,500 × *g* for 10 min and 40,000 × *g* for 120 min at 4°C. Pellets were resuspended in mito-membrane buffer and stored at -80°C. For BN-PAGE, unsealed mitochondrial membranes were mixed with solubilisation buffer (750 mM aminocaproic acid, 50 mM Bis-tris-HCl pH 7.0, 0.5 mM EDTA, 1 % (w/v) dodecyl maltoside) and incubated on ice for 5 min followed by centrifugation at 16,100 × g, at 4°C, for 10 min. The supernatant containing solubilized membrane proteins were diluted to 0.25 μg μl^-1^ protein (Bradford assay) with solubilization buffer and 0.25% (w/v) Coomassie G250. Typically, 6.25 μg protein was loaded per lane. 5 μl NativeMark^TM^ was used as a molecular weight marker. All BN-PAGE performed used NativePage^TM^ 4-16% Bis-Tris gels (Invitrogen), with 1 mm spacers. The anode buffer was 50 mM Bis-Tris-HCl, pH7.0, cathode buffer was 50 mM Tricine, 15 mM Bis-Tris-HCl, pH7.0. Gels were run in a XCell SureLock^TM^ system (Thermofisher) at 4°C, for the first 45 min at 100 V (max 10 mA) with cathode buffer containing 0.02% (w/v) Coomassie G250, then for ~2.5 h at 250 V (max 15 mA) with cathode buffer containing 0.002% (w/v) Coomassie G250, until the dye front exited the bottom.

BN gels or PVDF membrane were stained in Coomassie solution (45% (v/v) MeOH and 5% (v/v) acetic acid, 0.05% Coomassie R250) for 1-12 hours. Gels were destained using a solution of 45% (v/v) methanol and 5% (v/v) acetic acid.

### Protein blot analysis

Mitochondrial membranes were mixed with Laemmli buffer (2% [w/v] SDS, 125 mM Tris-HCl pH 6.8, 10% [w/v] glycerol, 0.04% [v/v] β-mercaptoethanol, and 0.002% [w/v] bromophenol blue) and separated on a 15% SDS-PAGE gel. Proteins were transferred under semi-dry conditions to nitrocellulose membrane (Protran^TM^). Equal loading and transfer was confirmed using Ponceau S stain. Proteins were labelled with antibodies and detected using secondary horseradish peroxidase-conjugated antibodies and chemiluminescence. Mouse monoclonal antibodies against the NUBM and NUCM subunit complex I were a gift from Volker Zickermann. Rabbit polyclonal antibodies against Sdh2 and Aco1 were raised against recombinant *Saccharomyces cerevisiae* proteins and previously reported (38); antibodies against *Y. lipolytica* Ind1 are described in (8).

Protein was transferred from the BN-PAGE gel to a PVDF membrane (0.2 micron, Millipore^TM^) using wet transfer and BN cathode buffer, for 600 minutes at 40 mA, (maximum voltage 100 V). After transfer immunolabelling was carried out as described above.

### NADH dehydrogenase assays

BN gels were stained for complex I, as described in (23). Briefly, BN gels were equilibrated in 0.1 M Tris-HCl pH 7.4, then incubated with new buffer containing 0.2 mM NADH and 0.1% (w/v) nitroblue tetrazolium. After the desired staining intensity was reached, the reaction was stopped using a solution of 45% (v/v) MeOH and 5% (v/v) acetic acid. Destaining using a solution of 45% (v/v) MeOH and 5% (v/v) acetic acid was performed for optimal enzyme stain visualisation.

NADH:HAR oxidoreductase activity was assayed as described (8). NADH:HAR oxidoreductase activity was measured as NADH oxidation in the presence of the artificial electron acceptor hexaammineruthenium(III) chloride (HAR). A 1 ml, 10 mm quartz cuvette was used and. Activity for 25 μg mitochondria was measured in a 1-ml volume of 20 mM HEPES-NaOH pH 8.0, 2 mM NaN_3_, 0.2 mM NADH, 2 mM HAR and 40 μg ml^-1^ alamethicin. Measurements made in a quartz cuvette using a UV/Vis spectrophotometer (Jasco, V-550) and started after addition of HAR.

dNADH:DBQ oxidoreductase activity was measured as dNADH oxidation activity in the presence of the ubiquinone analogue n-decylubiquinone (DBQ) as an electron acceptor. Activity for 50 μg mitochondria was measured in a 1-ml volume of 20 mM MOPS-NaOH pH 7.4, 50 mM NaCl, 2 mM KCN, 0.1 mM dNADH and 0.1 mM DBQ. Measurement started after addition of DBQ. 0.01 mM of the complex I inhibitor Piericidin A was added at the end of the reaction to ensure that dNADH:DBQ oxidoreductase activity is due to complex I.

## Acknowledgements

We thank Ulrich Brandt, Radboud Center for Mitochndrial Medicine, Nijmegen, the Netherlands, for yeast strains *nubm*Δ, *nucm*Δ, *nucm*Δ + *NUCM*^Y144F^, *nukmΔ;* and for antibodies again NUBM and NUCM; Holger Prokisch for sharing information on the NUBPL G287C variant.

This work was supported by the John Innes Foundation [studentship to A.E.M.]; the Spooner Girls Foundation [to V.E.K.], the UC Irvine Institute for Clinical and Translational Science and Center for Autism Research [to V.E.K.]; and the Biotechnology and Biological Sciences Research Council (BBSRC) [BB/J004561/1 to J.B.].

## Abbreviations

BN-PAGE: ,Blue native PAGE
CBB: , Coomassie Brilliant Blue
cWT: , complemented wild type
DBQ: , n-decylubiquinone
dNADH: , deamino-NADH
e.v.: , empty vector
FeS: , iron-sulfur
HAR: , hexaammineruthenium(III) chloride
Ind1 (protein) and IND1 (gene): , Iron-sulfur protein required for NADH dehydrogenase
MRI: , magnetic resonance imaging
NBT: , nitro-blue tetrazolium
NUBPL: , Nucleotide binding protein-like
PAGE: , polyacrylamide gel electrophoresis
ROS: , reactive oxygen species
RT-PCR: , reverse transcription - polymerase chain reaction

